# Brain white matter structure and language ability in preschool-aged children

**DOI:** 10.1101/184978

**Authors:** Matthew Walton, Deborah Dewey, Catherine Lebel

**Author notes:** Corresponding Author: Catherine Lebel, Room B4-513, Alberta Children’s Hospital, 2888 Shaganappi Trail NW, Calgary, Alberta, Canada, T3B 6A8.

## Abstract

Brain alterations are associated with reading and language difficulties in older children, but little research has investigated relationships between early language skills and brain white matter structure during the preschool period. We studied 68 children aged 3.0-5.6 years who underwent diffusion tensor imaging and participated in assessments of Phonological Processing and Speeded Naming. Tract-based spatial statistics and tractography revealed relationships between Phonological Processing and fractional anisotropy and mean diffusivity in bilateral ventral white matter pathways, the corpus callosum, and corticospinal tracts. The relationships observed in left ventral pathways are consistent with studies in older children, and demonstrate that structural markers for language difficulties are apparent as young as 3 years of age. Our findings in right hemisphere areas that are not as commonly found in adult studies suggest that young children rely on a widespread network for language processing that becomes more specialized with age.

## 1. Introduction

One of the most important developmental gains seen in young children is the acquisition of language skills. Language ability in early childhood has been associated with future reading success, and can affect academic achievement, mental health, and future career prospects (Carroll, Maughan, Goodman, & Meltzer, 2005). Although most children develop fluent reading skills, 5-17% will be diagnosed with dyslexia, a disorder characterized by reading problems that may persist throughout life (Shaywitz, 1998). Interventions in kindergarten-aged children at risk for dyslexia have been shown to be effective at improving grade school reading ability (Elbro & Petersen, 2004); however, reading disabilities are typically not diagnosed and treated until the third grade, when reading delays are clearly measureable (Gabrieli, 2009). There is general consensus that the roots of dyslexia begin before initial reading instruction, and that assessment of phonological processing skills could assist in early identification of children at risk for dyslexia (Gabrieli, 2009).

Neuroimaging studies have helped develop a better understanding of the relationship between white matter brain architecture and language and reading (Smits, Jiskoot, & Papma, 2014), and contributed to a dual stream model of language that includes dorsal and ventral pathways (Hickok & Poeppel, 2004). The dorsal pathway, specifically the arcuate fasciculus (AF), connects left frontal and temporal-parietal regions (Dick & Tremblay, 2012), is related to sensory-motor mapping of sound to articulation (Saur et al., 2008), and sustains phonological aspects of reading (Vandermosten, Boets, Poelmans, et al., 2012; Vigneau et al., 2006). The ventral pathways, which include the uncinate fasciculus (UF), inferior longitudinal fasciculus (ILF), and inferior fronto-occipital fasciculus (IFOF), connect frontal, temporal, and occipital regions (Dick & Tremblay, 2012), are involved with linguistic processing of sound to meaning (Saur et al., 2008), and are thought to sustain orthographic aspects of reading, such as the ability to identify words by sight (Jobard, Crivello, & Tzourio-Mazoyer, 2003; Vandermosten, Boets, Poelmans, et al., 2012).

Diffusion tensor imaging (DTI) studies in school-aged children and adults demonstrate that language and reading abilities have positive relationships with fractional anisotropy (FA) and negative relationships with mean diffusivity (MD) in left temporal parietal and frontal areas (Vandermosten, Boets, Wouters, & Ghesquiere, 2012). Furthermore, longitudinal studies show that gains in language abilities are associated with changes in white matter structure over time (Keller & Just, 2009; Yeatman et al., 2012). While these studies convincingly demonstrate a relationship between reading/language and brain structure in older children and adults, it is unclear whether these relationships are identifiable before children begin to read. Two DTI studies investigated language abilities in children in Kindergarten, aged 5-6 years, and showed positive relationships between phonological processing and FA in the left AF (Saygin et al., 2013), and in the bilateral AF and IFOF (Vandermosten et al., 2015). Another DTI study examined children as young as 5 years and found that pre-readers with a family history of dyslexia had lower FA in the temporal parietal segment of the AF, when compared to pre-readers with no family history of dyslexia (Wang et al., 2016). Similarly, Langer et al. (2015) found that infants aged 5-18 months with a family history of dyslexia had lower FA in the AF compared to infants with no family history, and that expressive language ability correlated with FA in the left AF across groups. Relationships between receptive and expressive language ability and myelin water fraction have also been observed in children 1-6 years of age (O’Muircheartaigh et al., 2013; O’Muircheartaigh et al., 2014). The above mentioned studies show relationships between white matter structure and reading/language skills are apparent at a young age; however, no studies have yet examined the relationships between white matter structure and specific pre-reading skills such as phonological processing and speeded naming in children younger than 5 years. Understanding the relationships between brain structure and language abilities predictive of future reading skill is necessary to characterize the neurological basis of language disorders and the roots of reading difficulties. Therefore, the goal of this study was to investigate the relationship between white matter structure and pre-reading skills in a large group of typically developing preschool aged children.

## 2 Methods

### 2.1 Participants

Sixty-eight children aged 3.0-5.6 years (4.0 ± 0.6 years, 31 females and 37 males) were recruited from an ongoing prospective study that recruited women during pregnancy (Kaplan et al., 2014). All children were healthy, free from neurological or developmental disorders, had no history of head trauma and had no contraindications to MRI scanning (e.g., metal implants). Gestational age at birth ranged from 36.7-41.9 weeks (39.5 ± 1.7 weeks). All children spoke English as a primary language, with 9 children identified as coming from a bilingual home. Family history of reading disability was collected from parental reports, which identified 7 children with a first or second degree relative that had been diagnosed with developmental dyslexia. Children’s hand preference was assessed by parental questionnaire, of which 7 participants were identified as being predominately left handed, with the remaining 61 being predominately right handed (no participants were identified as ambidextrous). Number of years of maternal post-secondary education was collected as a proxy for socio-economic status, and ranged from 1-12 years, with a mean of 5.0 ± 2.8 years. The institutional ethics review board approved this study, and informed consent was obtained from participant’s legal guardian(s), and verbal assent was obtained from the children.

### 2.2 Language Assessments

All participants completed the Phonological Processing and Speeded Naming subtests of the NEPSY-II (Korkman, Kirk, & Kemp, 2007). The Phonological Processing test assesses phonemic awareness, whereas the Speeded Naming subtest measures rapid semantic access to and production of names of colors. Phonological Processing Scaled Scores and Speeded Naming Combined Scaled Scores were used for the analysis. The Speeded Naming Combined Scaled Score takes into account both the speed and accuracy of the answers given by the child. These language assessments took approximately 10 minutes to complete, and were administered on the same day as MRI scanning.

### 2.3 MRI Scanning

Imaging took place at the Alberta Children’s Hospital on the same General Electric 3T MR750w system using a 32-channel head coil (GE, Waukesha, WI). Children were scanned while awake and watching a movie, or while sleeping without sedation. Whole-brain diffusion weighted images were acquired using a single shot spin echo echo-planar imaging protocol, with 30 gradient encoding directions at b=750s/mm^2^ and 5 images without gradient encoding (b=0s/mm^2^). The DTI sequence was acquired with the following parameters: TR = 6750 ms, TE = 79 ms, 50 axial slices with a 2.2mm thickness (no gap), 1.6x1.6x2.2mm^3^ resolution, anterior-posterior phase encoding, scan time = 4:03 minutes.

### 2.4 Data processing

DTI data was quality checked and processed through in house, Matlab-based software designed to detect and remove motion-corrupted volumes. Motion in the DTI images was quantified based on the presence of black lines in the dataset. A Prewitt horizontal edge-enhancing filter was applied to sagittal and coronal views of the DTI data, then a Hough transform was used to detect and quantify black lines in these views. If the quantity of black lines exceeded a tested threshold, the volume was labeled as motion corrupted and removed from the data set. From an original sample of 74 children, 6 participants were excluded from the study because they had 15 or more diffusion weighted volumes removed for motion corruption. Of the 68 children included in the final sample, 97% of included participants had fewer than 5 volumes removed, and 76% of children had no motion-corrupted volumes. The number of motion-corrupted volumes was not significantly correlated with age (p=0.31) or language scores (Phonological Processing p=0.70; Speeded Naming p=0.37).

Data was processed using FSL’s diffusion pipeline (Jenkinson, Beckmann, Behrens, Woolrich, & Smith, 2012), including eddy current correction and simple head motion correction using an affine registration to a reference volume, fitting of a diffusion tensor model at each voxel, and calculation of fractional anisotropy (FA), mean diffusivity (MD), axial diffusivity (AD), and radial diffusivity (RD). All processed images were visually inspected to ensure high quality.

### 2.5 Data analysis

Whole-brain voxel-wise analysis was performed using FSL’s Tract Based Spatial Statistics (TBSS) (Smith et al., 2006). Using non-linear registration, each participant’s FA image was aligned to that of every other participant. The most representative participant (i.e., the one that required the least warping for all other participants) was chosen as the target image. The target image was then aligned into MNI152 standard space using affine registration, and every image was transformed into 1x1x1mm MNI152 space by combining the nonlinear transform to the target FA image with the affine transform from that target to MNI152 space. This method was selected based on the young age of the participants, which would not necessarily align correctly to the adult template provided in TBSS.

A mean white matter tract skeleton was created using an FA threshold of 0.3, then participants’ FA data was projected onto the skeleton and analyzed using voxel wise cross-subject statistics using non-parametric permutation testing (RANDOMISE). Significant clusters were identified at p<0.05 using threshold free cluster enhancement and Gaussian random field based family wise error rate correction for multiple comparisons (Smith and Nichols, 2009). Clusters were labeled using the NatBrainLab white matter atlas (Thiebaut de Schotten et al., 2011). Projection onto the skeleton and statistics were repeated for MD, AD, and RD. Each DTI parameter was independently correlated with age-standardized Phonological Processing and Speeded Naming scores. Age, sex, hand preference, bilingualism, gestational age, family history of dyslexia, and maternal years of post-secondary education were included as covariates in the TBSS analysis.

To corroborate findings from the whole brain analysis, a follow up tractography analysis was performed. Identified tracts (from the NatBrainLab white matter atlas) containing significant clusters from the TBSS analysis were delineated using deterministic tractography in Trackvis (Wang, Benner, & Sorensen, 2007). Regions of interest were drawn on the most representative subject from the TBSS analysis, and warped to each subject’s native DTI space for automated tractography. Tracts were generated using an angle threshold of 30°, and quality checked manually. Approximately 15% of participants required small manual corrections to remove spurious fibers from the tract. Mean values of FA, MD, AD and RD were extracted for each tract and using a general linear model, relationships were examined between diffusion parameters and Phonological Processing and Speeded Naming standard scores, with age, sex, hand preference, bilingualism, gestational age, family history of dyslexia, and maternal years of post-secondary education entered as covariates.

We conducted a follow-up analysis to determine whether results were due to overall white matter development. Each participant’s average FA (and MD, RD, and AD) was calculated across the white matter skeleton, and included as a covariate in a supplementary analysis for both TBSS and tractography, along with the original covariates listed above.

## 3. Results

### 3.1 Language Assessments

On the Phonological Processing subtest, children achieved a mean aged-standardized scaled score of 11.2 ± 2.9 (range 3-17). For Speeded Naming, the mean scaled score was 11.5 ± 2.6 (range 5-16). Compared to the population norm of 10 ± 3, mean scores in our sample were significantly higher for both Phonological Processing (t=3.446, p=0.001) and Speeded Naming (t=4.824, p<0.0001), though still within the average range and spanning a wide range of abilities.

### 3.2 Brain-Language Relationships

TBSS analysis showed a significant positive relationship (p<0.05, family wise error rate corrected) between Phonological Processing scaled scores and FA, with significant clusters located in the left UF, IFOF and corticospinal tract (CST), bilateral internal capsule, and the genu and body of the corpus callosum (Figure 1A). No significant negative relationships were found between Phonological Processing scaled scores and FA.

A significant negative relationship was also seen between Phonological Processing scores and MD, with clusters in bilateral IFOF, left UF, and right ILF and posterior segment of the AF (Figure 1B). No significant positive relationships were found between Phonological Processing scaled scores and MD. No significant relationships were found between Phonological Processing and RD and AD, nor were any significant relationships found between Speeded Naming and any of the DTI measures.

**Figure 1.**
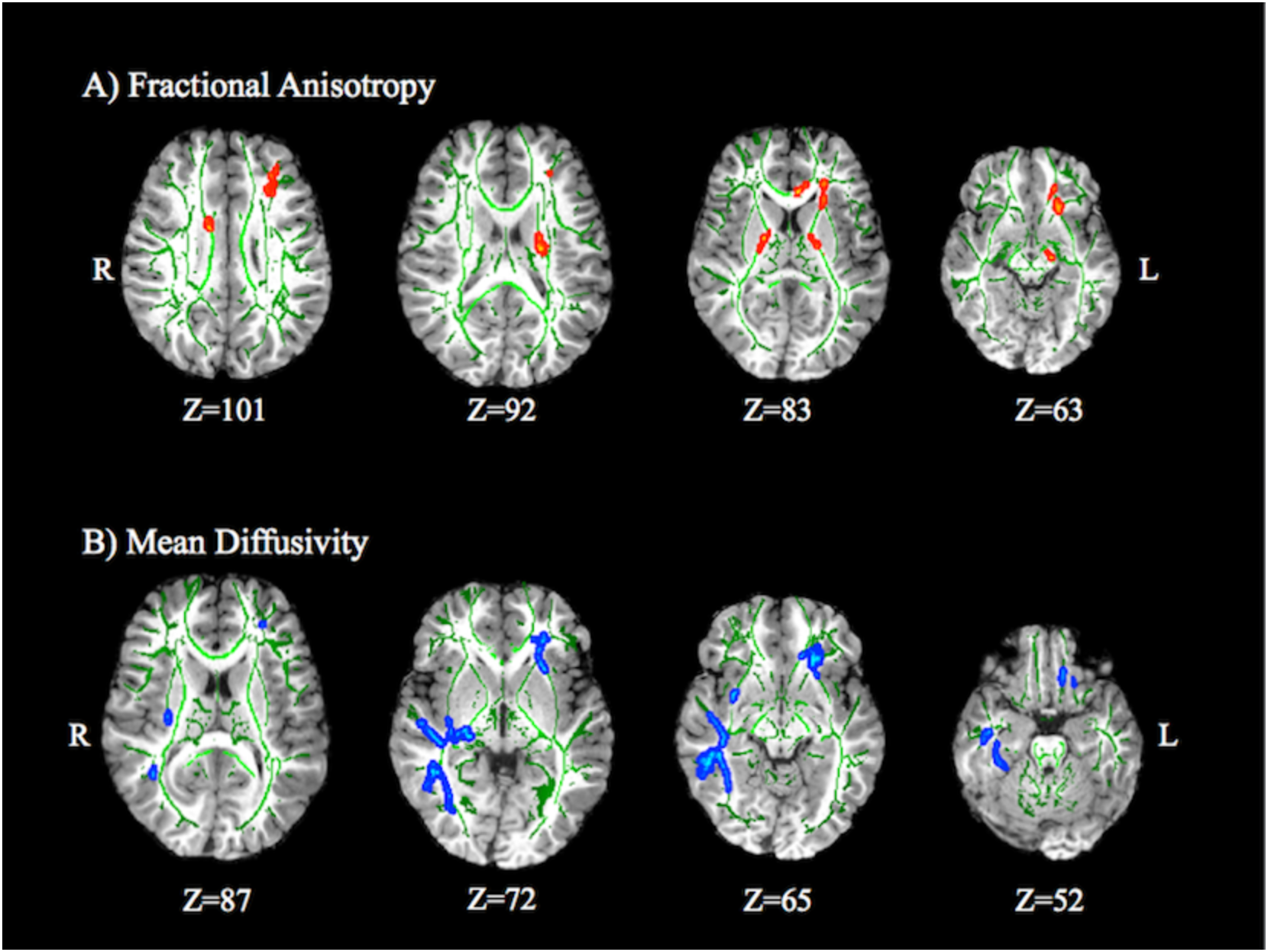
Tract based spatial statistics analysis of Phonological Processing scores and diffusion parameters. Significant relationships were found between Phonological Processing scores and (A) fractional anisotropy and (B) mean diffusivity in multiple white matter pathways. The white matter skeleton is shown in green, overlaid on a representative T1 map. Red areas represent a significant positive relationship, while blue areas indicate a significant negative relationship (p<0.05, family wise error rate controlled). For display purposes, significant clusters have been expanded from the skeleton to the full white matter space. For fractional anisotropy, significant positive relationships were observed in the left uncinate fasciculus (UF), inferior fronto-occipital fasciculus (IFOF), corticospinal tract (CST), bilateral internal capsule (IC), and body (BCC) and genu of the corpus callosum. For mean diffusivity, significant negative relationships were observed in bilateral inferior fronto-occipital fasciculi, left uncinate fasciculus, and right inferior longitudinal fasciculus and posterior segment of the arcuate fasciculus (AFp). No significant correlations were found between radial or axial diffusivity and Phonological Processing, nor were any relationships found between Speeded Naming and any of the DTI measures.

### 3.3 Tractography

The left UF and CST, bilateral IFOF, right AF and ILF, and genu and body of the corpus callosum were successfully isolated for each subject using tractography, with the exception of 3 participants in whom we were not able to delineate the right AF. We found a positive relationship (p ≤ 0.05) between Phonological Processing scores and FA in the bilateral IFOF, left CST, and the body and genu of the corpus callosum (Table 1, Figure 2A). There was a negative relationship (p ≤ 0.05) between Phonological Processing scores and MD and RD in the left IFOF, and UF, left CST, and genu of the corpus callosum (Table 1, Figure 2B). After Bonferroni correction for 8 comparisons (p<0.0063), the correlations between Phonological Processing scores and FA in the left IFOF, MD in the genu, and RD in the left IFOF, splenium, and genu remained significant. No relationships were found between Phonological Processing scores and AD, and between Speeded Naming and any DTI parameters.

**Table 1.**
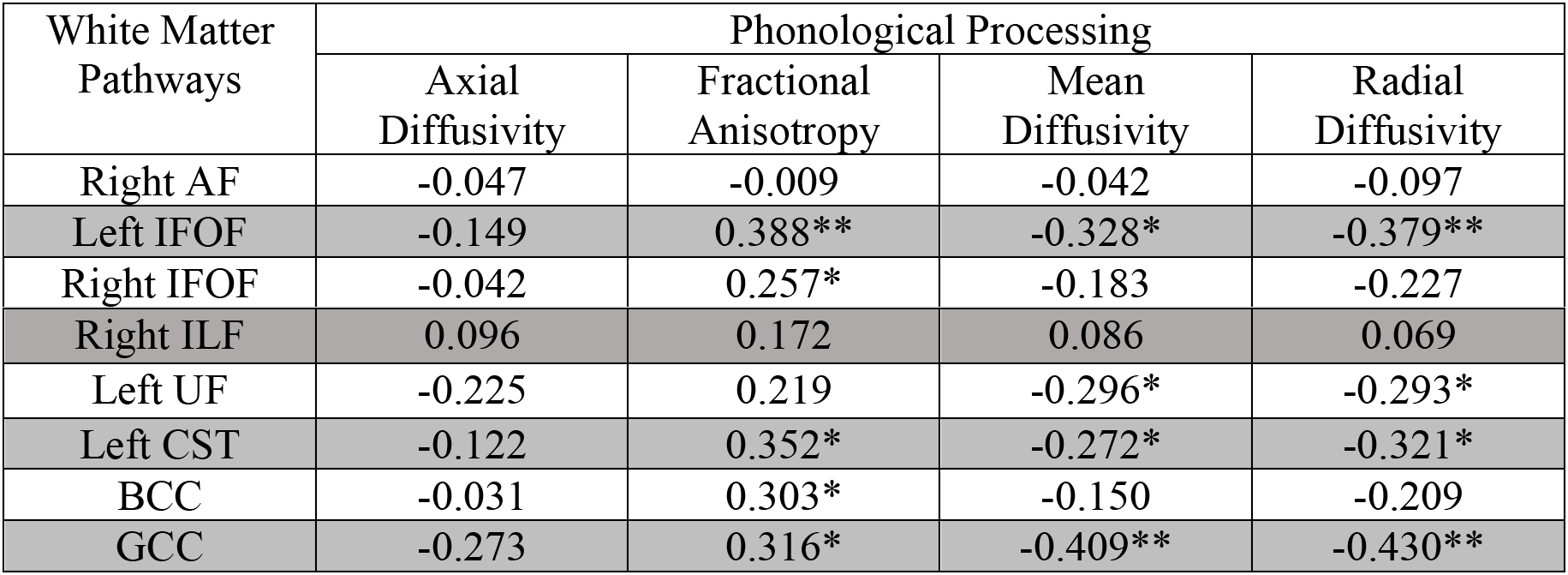
Relationships between language scores and tractography derived white matter tracts. Tractography was used to delineate pathways that had a significant finding in the TBSS analysis; Pearson r values are shown here. *p<0.05, **p<0.0063 (Bonferroni corrected for 8 tracts), Abbreviations: AF (arcuate fasciculus), IFOF (inferior fronto-occipital fasciculus), ILF (inferior longitudinal fasciculus), UF (uncinate fasciculus), CST (corticospinal tract), BCC (body of corpus callosum), GCC (genu of corpus callosum).

| White Matter Pathways | Phonological Processing |  |  |  |
| --- | --- | --- | --- | --- |
|  | Axial Diffusivity | Fractional Anisotropy | Mean Diffusivity | Radial Diffusivity |
| Right AF | -0.047 | -0.009 | -0.042 | -0.097 |
| Left IFOF | -0.149 | 0.388** | -0.328* | -0.379** |
| Right IFOF | -0.042 | 0.257* | -0.183 | -0.227 |
| Right ILF | 0.096 | 0.172 | 0.086 | 0.069 |
| Left UF | -0.225 | 0.219 | -0.296* | -0.293* |
| Left CST | -0.122 | 0.352* | -0.272* | -0.321* |
| BCC | -0.031 | 0.303* | -0.150 | -0.209 |
| GCC | -0.273 | 0.316* | -0.409** | -0.430** |

**Figure 2.**
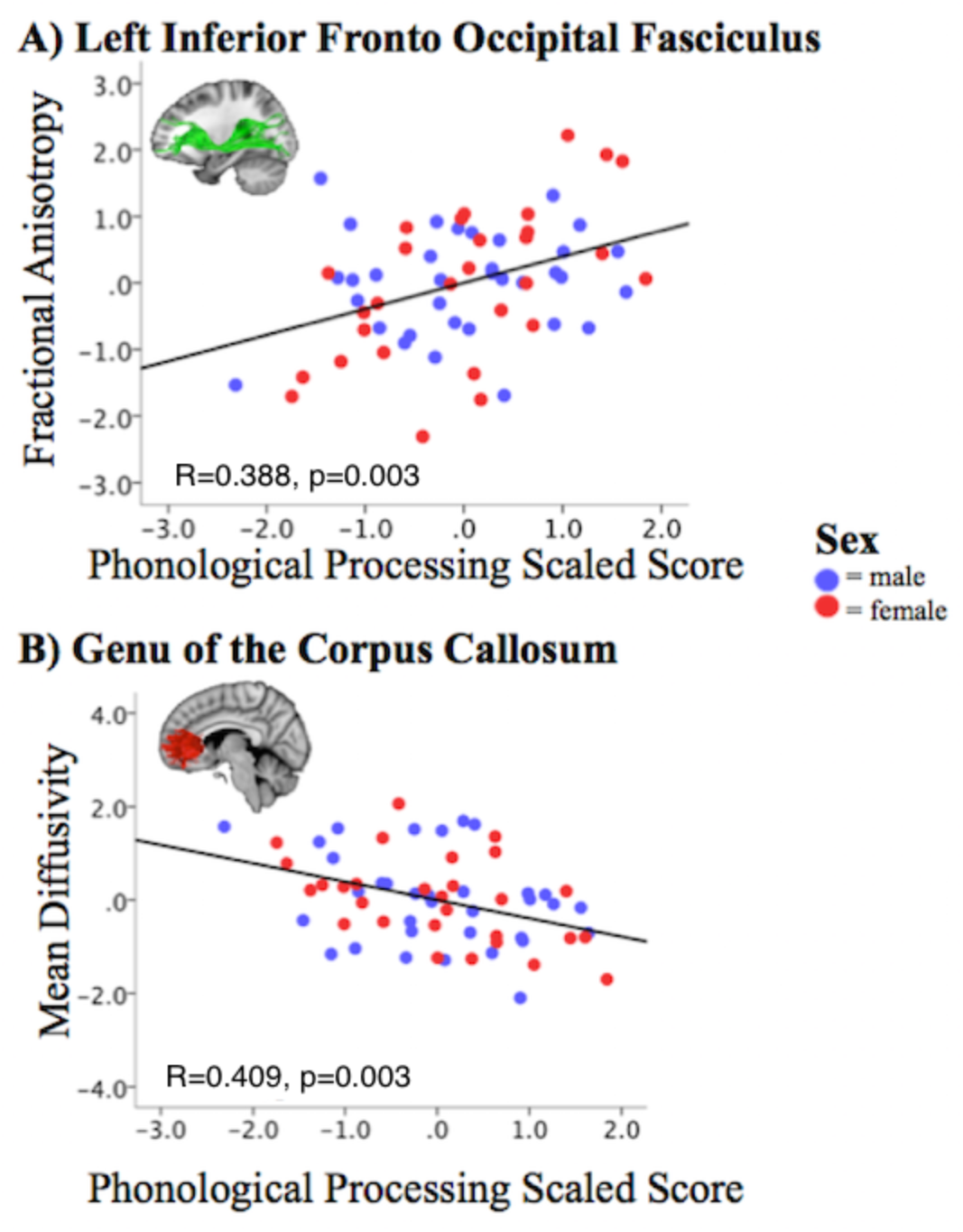
Relationship between DTI parameters from tractography derived pathways and Phonological Processing. Scatterplots show the relationships between Phonological Processing Scaled scores and DTI measurements for two sample regions: fractional anisotropy of the left inferior fronto-occipital fasciculus (A) and mean diffusivity of the genu of the corpus callosum (B). Males are marked in blue and females in red. Values shown on graphs are standardized residuals. Relationships between phonological processing and fractional anisotropy, mean diffusivity, and radial diffusivity were also noted in other pathways (see Table 1).

### 3.4 Sex Differences

Females scored slightly higher than males on both Phonological Processing (female=11.3, male=11.1) and Speeded Naming (female=12.1, male=11.1); however, these differences were not significant in an independent samples t-test. There were also no significant sex differences on any of the diffusion measures assessed using TBSS or tractography.

### 3.5 General White Matter Development

When controlling for average FA, the TBSS results remained largely the same, with significant correlations between Phonological Processing and FA in the uncinate and IFOF, internal capsule, and genu of the corpus callosum (Supplementary Figure 1). Some areas of the CST and body of the corpus callosum did not meet significance thresholds in the TBSS analysis when controlling for average FA. TBSS results for MD remained almost identical when controlling for average MD, and no significant findings were noted in TBSS for AD or RD.

The tractography results indicated that all correlations remained significant when controlling for average diffusion values, including correlations between Phonological Processing and FA in the CST and BCC (Supplementary Table 1, Supplementary Figure 2). Pearson r values decreased in magnitude, however, and the correlations between Phonological Processing and FA in the left IFOF and MD in the genu of the corpus callosum were no longer significant after Bonferroni correction for multiple comparisons.

## 4. Discussion

In preschool-aged children, we showed that stronger phonological processing ability, which is predictive of future reading ability, is associated with white matter structure in several brain areas. Our findings of lower FA and higher MD in preschool children with weaker language ability are in agreement with DTI studies in older children and adults with language and reading disorders, and suggest that the brain systems underlying reading are structurally different in children with poor language abilities well before formal reading instruction begins. Because FA increases and MD decreases in early childhood (Hagmann et al., 2010; Hermoye et al., 2006; Krogsrud et al., 2015), our results may indicate that children with stronger language skills display a more mature arrangement of brain structure. Most relationships remained significant when controlling for average diffusion parameters, suggesting that the results reflect specific properties of the tracts identified rather than overall white matter maturation.

Our results demonstrate relationships between phonological ability and white matter structure in bilateral ventral language pathways. Although it is not possible to fully distinguish tracts based on the significant clusters identified in TBSS, our follow-up tractography analysis strongly suggests the inferior fronto-occipital fasciculus is involved. Previous DTI studies on older children and adults have typically associated the dorsal white matter system (arcuate fasciculus) with phonological ability (Saygin et al., 2013; Vandermosten et al., 2015; Nagy, Westerberg, & Klingberg, 2004; Yeatman et al., 2011; Gold, Powell, Xuan, Jiang, & Hardy, 2007; Nagy et al., 2004; Steinbrink et al., 2008; Vandermosten, Boets, Poelmans, et al., 2012). Findings have been mixed for ventral white matter pathways, with some studies reporting significant language-white matter structure relationships (Odegard, Farris, Ring, McColl, & Black, 2009; Rimrodt, Peterson, Denckla, Kaufmann, & Cutting, 2010; Steinbrink et al., 2008; Vandermosten, Boets, Poelmans, et al., 2012; Vandermosten et al., 2015), and others failing to find correlations (Beaulieu et al., 2005; Deutsch et al., 2005; Dougherty et al., 2007; Klingberg et al., 2000; Nagy et al., 2004; Niogi & McCandliss, 2006; Saygin et al., 2013). In addition to the dorsal and ventral pathways, recent studies have suggested shorter frontal lobe connections such as the frontal aslant and fronto-orbital-polar tract may be associated with language ability (Catani et al., 2012). The frontal aslant tract, which connects Broca’s area with anterior cingulate and pre-supplementary motors areas was shown to be left-lateralized in a group of healthy adults (Catani et al., 2012). Furthermore, the length of the left frontal aslant tract was associated with receptive language in 5-8 year olds, suggesting that this pathway may be important for language development (Broce et al., 2015). Our TBSS results also appear to implicate these tracts in pre-reading skills, as phonological processing was correlated with FA and MD in a region that corresponds to the left frontal aslant tract. However, we were not able to isolate the frontal-aslant tract using tractography, so this finding remains speculative.

Consistent with our findings, Vandermosten et al. (2015) reported a positive relationship between FA in the bilateral IFOF and phonological ability in 71 pre-readers (mean age 5.1Y), and suggested that because dorsal language pathways mature later than ventral ones (Brauer, Anwander, & Friederici, 2011; Giorgio et al., 2008; Lebel et al, 2008; Lebel & Beaulieu, 2011), younger children may rely more heavily on ventral pathways for phonological processing than older children. Although our results do not relate specifically to reading difficulty/disorders, our findings are in line with previous studies that have shown that pre-readers as young as 5 years (Wang et al., 2016) and infants (Langer et al., 2015) with a family history of dyslexia have lower FA than prereaders/infants with no family history of dyslexia.

We failed to find a significant relationship between speeded naming ability and any of the DTI parameters. This is consistent with one study on pre-readers, which also failed to find significant correlations between FA and speeded naming (Saygin et al., 2013), but contrasts another study (Vandermosten et al., 2015), which found a positive relationship between FA of the right IFOF and speeded naming ability in pre-readers aged 5 years. The lower mean age in our study may indicate that relationships between speeded naming ability and white matter brain structure do not appear until slightly later in childhood. Alternatively, participant scores for Speeded Naming had a narrower range than Phonological Processing, reducing the statistical power to detect a relationship.

As expected, we observed many relationships between language and white matter pathways in the left hemisphere; however, there were also several significant associations between language and right hemisphere pathways. Previous functional (Raschle, Zuk, & Gaab, 2012; Turkeltaub, Gareau, Flowers, Zeffiro, & Eden, 2003; Yamada et al., 2011) and structural imaging (O’Muircheartaigh et al., 2013; Raschle, Chang, & Gaab, 2011; Vandermosten et al., 2015) studies suggest that the right hemisphere is involved in language processing, particularly in children younger than 6 years (O’Muircheartaigh et al., 2013; Raschle et al., 2011; Raschle et al., 2012; Vandermosten et al., 2015; Yamada et al., 2011). Functional imaging studies show that adult-like left hemisphere language lateralization emerges around age 5 years (Weiss-Croft & Baldeweg, 2015) from an earlier bilateral network (Friederici, Brauer, & Lohmann, 2011; Kadis et al., 2011; Ressel, Wilke, Lidzba, Lutzenberger, & Krageloh-Mann, 2008; Yamada et al., 2011); DTI studies confirm structural leftward-lateralization in most older children and adults (Lebel & Beaulieu, 2011; Qiu et al., 2011). The only previous study examining the relationship between white matter and language abilities in healthy preschoolers showed relationships between myelin water fraction asymmetry and expressive and receptive language that changed with age, but stabilized around 4 years (O’Muircheartaigh et al., 2013). Thus, our current results may represent the transition from a bilateral language network to one in which language is more left-lateralized. Handedness is not well established until age 6 years (Bryden et al., 2000), and is not associated with structural lateralization of language areas in older children and adults (Vernooij et al., 2007). Nonetheless, to ensure it was not driving our results, it was included as a covariate in all analyses, and results were similar when left-handers were excluded.

In addition to ventral white matter pathways, we found language abilities were related to the structure of less traditional language or reading areas including the corpus callosum and corticospinal tracts. Some previous DTI studies examining reading ability found that lower FA in the splenium of the corpus callosum was related to better reading ability (Dougherty et al., 2007; Frye et al., 2008; Odegard et al., 2009), whereas others indicate that higher FA or lower MD are associated with stronger reading skills (Lebel et al., 2013; Qiu, Tan, Zhou, & Khong, 2008). Our results are congruent with the latter, as we found higher FA and lower MD in the corpus callosum were related to higher Phonological Processing scores. We also observed significant relationships between Phonological Processing scores and FA and in the left corticospinal tract. This result supports the idea that language function in the brain becomes more specialized over time, as brain structure-language associations in the corticospinal tracts are not as apparent in older children or adults (Deutsch et al., 2005; Gold et al., 2007).

The results from our tractography analysis show that language relationships with FA and MD values appear to be driven by lower RD in children with better language ability. Animal studies suggest that AD is more related to axonal properties such as axonal diameter (Song et al., 2003), while RD is more related to myelination and axonal packing (Beaulieu, 2002; Song et al., 2002); MD varies with both axonal properties and myelination. Thus, lower RD in children with stronger language abilities, coupled with the lack of AD findings, may suggest differences in myelination or axonal packing. Studies in older children and adults have more consistently found relationships between reading ability and RD than AD (Dougherty et al., 2007; Keller & Just, 2009; Vandermosten, Boets, Poelmans, et al., 2012; Yeatman et al., 2011), which might indicate that language-white matter relationships are influenced by myelination throughout development. However, given the discrepancy between the size of individual axons and the voxel size of DTI data, any interpretation at a cellular level should be made with caution. Future work using more advanced imaging techniques for myelination may be able to more specifically characterize the biological processes underlying these relationships. However, scanning times must remain short to be feasible with young, non-sedated children.

Longitudinal studies are necessary to provide more information about relationships between developmental trajectories in brain structure and language abilities in young children. A longitudinal study in children 7-12 years old demonstrated different trajectories of FA values in reading tracts between good readers and poor readers (Yeatman et al., 2012). The results showed that children with better phonological awareness had lower initial FA values that increased over time, suggesting that young children with better phonological skills may have lower FA than other children. However, we found higher FA and lower MD in children with better phonological processing abilities. It remains unclear how the trajectory of brain development in early childhood is associated with pre-reading skills.

An advantage of this study is the combined TBSS and tractography analyses, which showed good consistency. By performing the whole-brain TBSS analysis first, we were able to forgo a priori anatomical hypotheses, which allowed us to detect relationships between language ability and white matter structure across the brain with good sensitivity to small, local changes. The confirmation of our findings by tractography suggests that the changes are not associated with only small areas of each tract. While our tractography method calculated an average diffusion parameter value across the whole tract, newer methods now allow for statistical analyses along individual segments of the tract. Future work using along-tract statistical methods may provide more information to language-white matter relationships. A limitation of this study is the cross-sectional design, which precludes measurement of brain development over time, and means that we can only infer that higher FA and lower MD represents a more mature brain structure in children with stronger language abilities. Future studies with longitudinal data are required to determine how trajectories of brain development in children vary with language ability. The lack of general cognitive data means that we cannot determine whether the relationships are specific to language abilities or related to other cognitive functions (e.g., visuospatial skills, attention, general intelligence). While our study attempted to recruit a sample of children with a wide range of language abilities, the scores of the children were slightly higher than average. This makes interpreting our results in the context of a normal population more difficult.

In conclusion, using DTI, we identified relationships between white matter structure and pre-reading language abilities in preschool-aged children. Both TBSS analysis and tractography showed that higher FA and lower MD were associated with better performance on the Phonological Processing subtest of the NEPSY-II, a measure which is predictive of future reading ability. Given that white matter FA increases, and MD decreases over time in children, our results suggest that young children with stronger language ability have more mature brain structure. Our results show that associations between brain structure and language exist before formal reading instruction has begun, and support the theory that altered brain structure may be a cause of poor reading, as opposed to a consequence of it. Ultimately, these results could assist in developing and optimizing interventions for children with language and reading difficulties.

## Acknowledgements

This work was supported by the Canadian Institutes of Health Research (CIHR) (funding reference numbers IHD-134090, MOP-136797, New Investigator Award to CL), and the NSERC CREATE International and Industrial Imaging Training (I3T) Program (MW). The authors thank members of the APrON study for assistance with recruitment.

---

**Supplementary Table 1.**
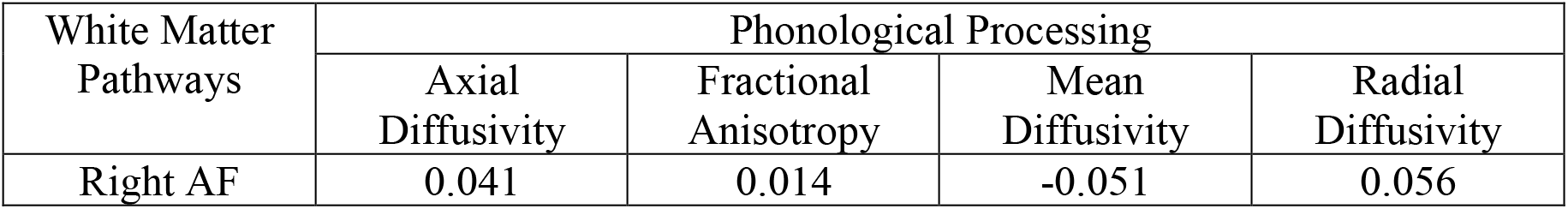
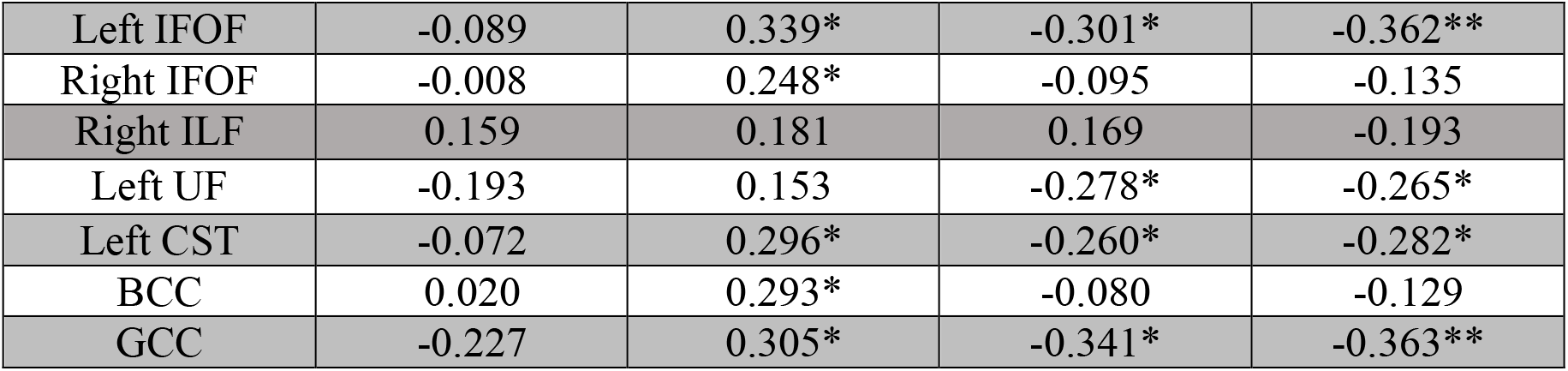
Relationships between language scores and tractography derived white matter tracts, controlling for general white matter development. R values are shown here for correlations between diffusion parameters and Phonological Processing Scores, after controlling for the average diffusion values across the white matter skeleton. *p<0.05, **p<0.0063 (Bonferroni corrected for 8 tracts), Abbreviations: AF (arcuate fasciculus), IFOF (inferior fronto-occipital fasciculus), ILF (inferior longitudinal fasciculus), UF (uncinate fasciculus), CST (corticospinal tract), BCC (body of corpus callosum), GCC (genu of corpus callosum).

| White Matter Pathways | Phonological Processing |  |  |  |
| --- | --- | --- | --- | --- |
|  | Axial Diffusivity | Fractional Anisotropy | Mean Diffusivity | Radial Diffusivity |
| Right AF | 0.041 | 0.014 | -0.051 | 0.056 |
| Left IFOF | -0.089 | 0.339* | -0.301* | -0.362** |
| Right IFOF | -0.008 | 0.248* | -0.095 | -0.135 |
| Right ILF | 0.159 | 0.181 | 0.169 | -0.193 |
| Left UF | -0.193 | 0.153 | -0.278* | -0.265* |
| Left CST | -0.072 | 0.296* | -0.260* | -0.282* |
| BCC | 0.020 | 0.293* | -0.080 | -0.129 |
| GCC | -0.227 | 0.305* | -0.341* | -0.363** |

**Supplementary Figure 1.**
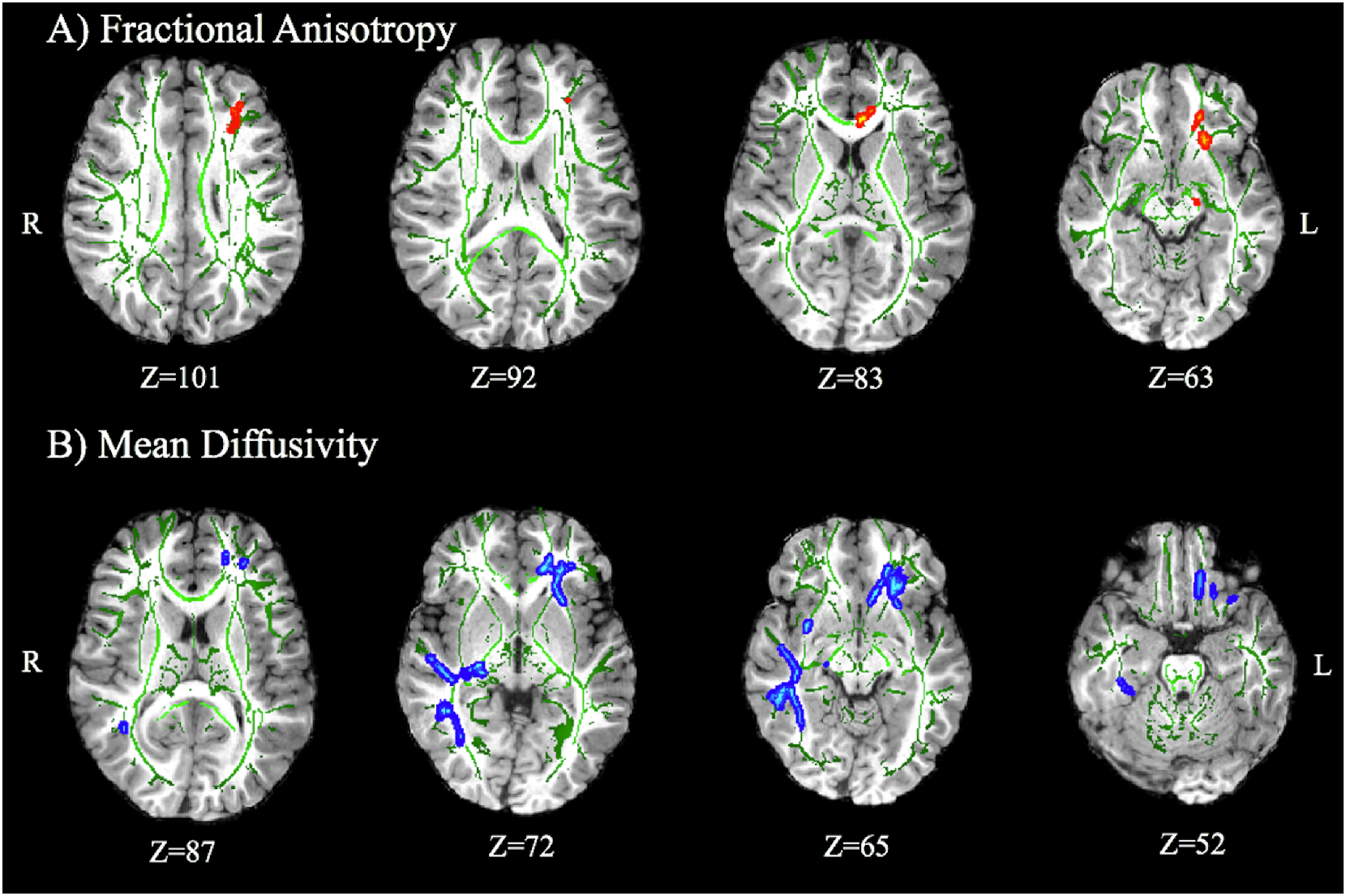
Tract based spatial statistics analysis of Phonological Processing scores and diffusion parameters, controlling for general white matter development. TBSS results while controlling for general white matter development (as measured by average diffusion parameters across the white matter skeleton) are similar to findings shown in Figure 1. For fractional anisotropy, some sections of the body of the corpus callosum and the corticospinal tracts no longer had significant relationships with Phonological Processing scores. For mean diffusivity, the result remained almost identical to those when not controlling for average MD. No significant correlations were found between radial or axial diffusivity and Phonological Processing, nor were any relationships found between Speeded Naming and any of the DTI measures.

**Supplementary Figure 2.**
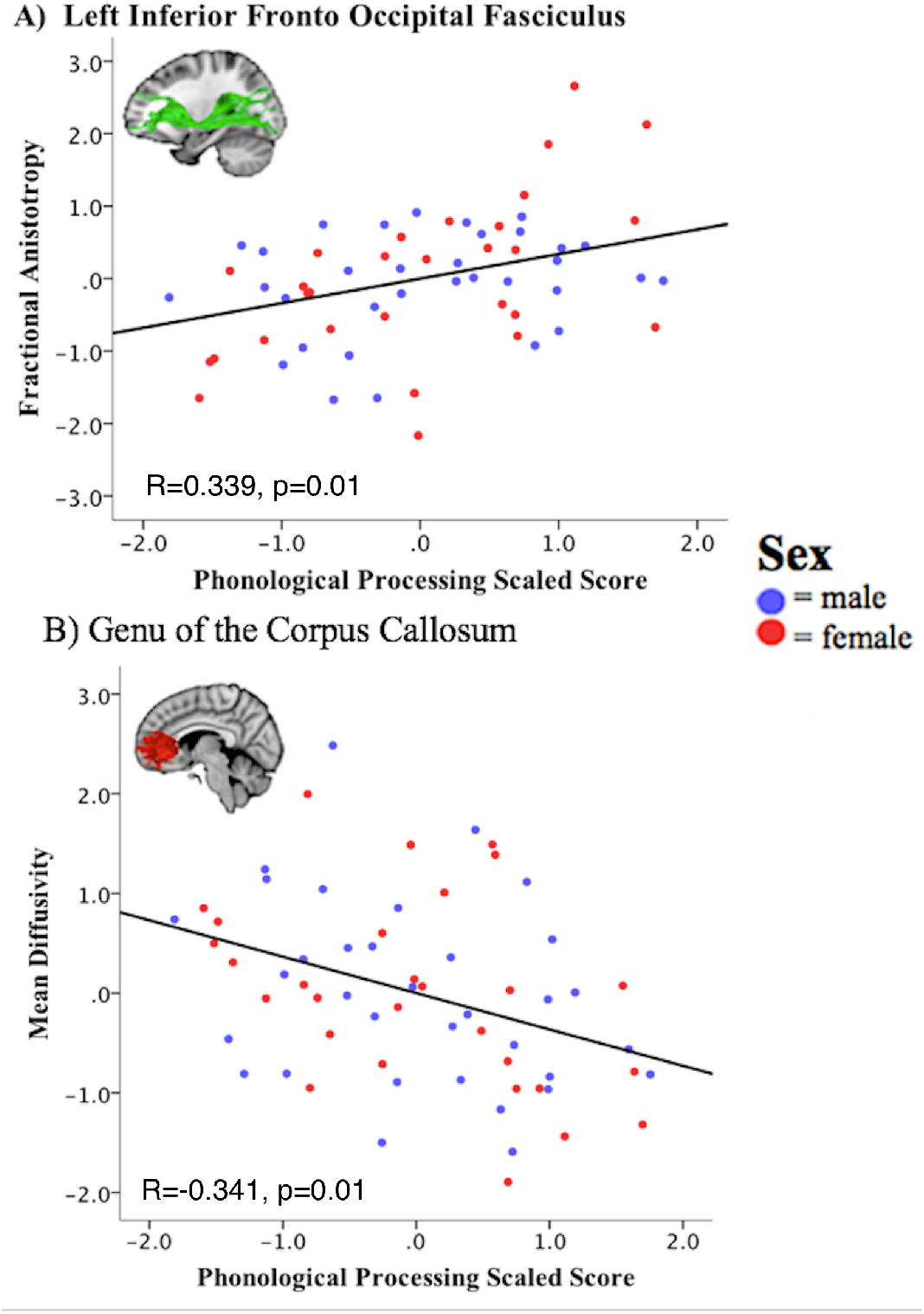
Relationship between DTI parameters from tractography derived pathways and Phonological Processing. Scatterplots show the relationships between Phonological Processing Scaled scores and fractional anisotropy (FA) of the left inferior fronto-occipital fasciculus (A), and mean diffusivity (MD) of the genu of the corpus callosum (B), after controlling for average FA and MD, respectively. Values shown on graphs are standardized residuals. Correlations decreased in magnitude after controlling for general white matter development, but were still significant in these tracts, and all others listed in Table 1.

